## Supplementary material for "FABP4-mediated lipid accumulation and lipolysis in tumor associated macrophages promote breast cancer metastasis": n/a

S-Figure 1

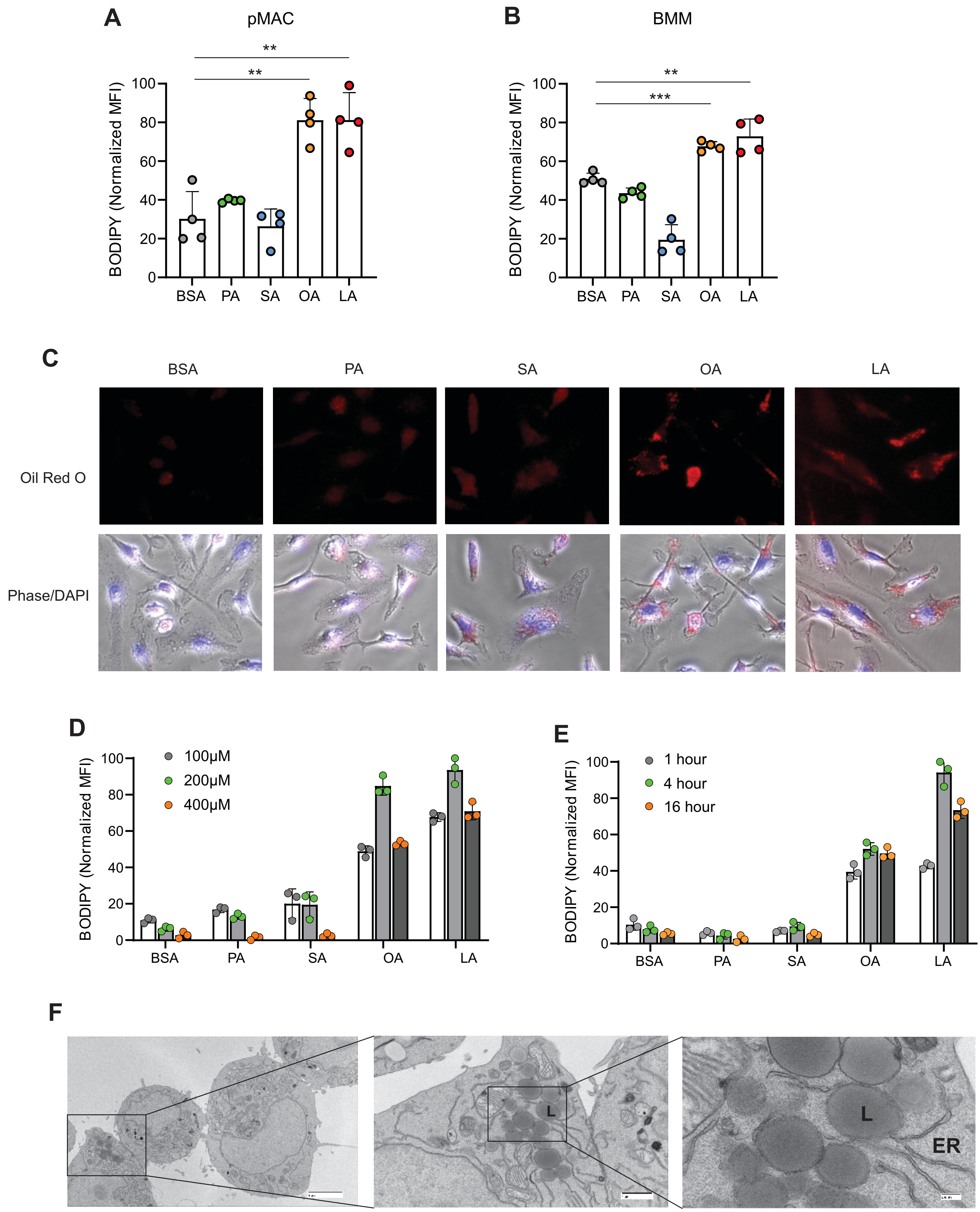

A
pMAC
**
**
BODIPY (Normalized MFI)
100
80
60
40
20
0
BSA
PA
SA
OA
LA
B
BMM
**
***
C
BSA
PA
SA
OA
LA
Oil Red O
Phase/DAPI
D
100μM
200μM
400μM
E
1 hour
4 hour
16 hour
F
L
L
ER

**Supplementary Figure 1 Unsaturated FA form lipid droplets in different macrophages.**

A, Flow cytometric analysis of lipid droplet formation in peritoneal macrophages treated with BSA and indicated dietary FA (100μM).

B, Flow cytometric analysis of lipid droplet formation in bone marrow-derived macrophages (BMM) treated with BSA and indicated dietary FA (100μM).

C, Confocal microscopy analysis of Oil Red O staining in BMM treated with 200μM of PA, SA, OA, LA and control BSA for 4h.

D, Titration of different indicated concentrations of dietary FA in forming lipid droplets in macrophage cell lines.

E, Measurement of lipid droplet formation in macrophages treated with indicated time periods of different dietary FAs (100μM).

F, Transmission electron microscope (TEM) images showing that LA-induced lipid droplets were formed in endoplasmic reticulum (ER) in macrophages.

Data are shown as mean ± SD in panel (** $p \le 0.01$, *** $p \le 0.001$, unpaired Student t test).

### S-Figure 2

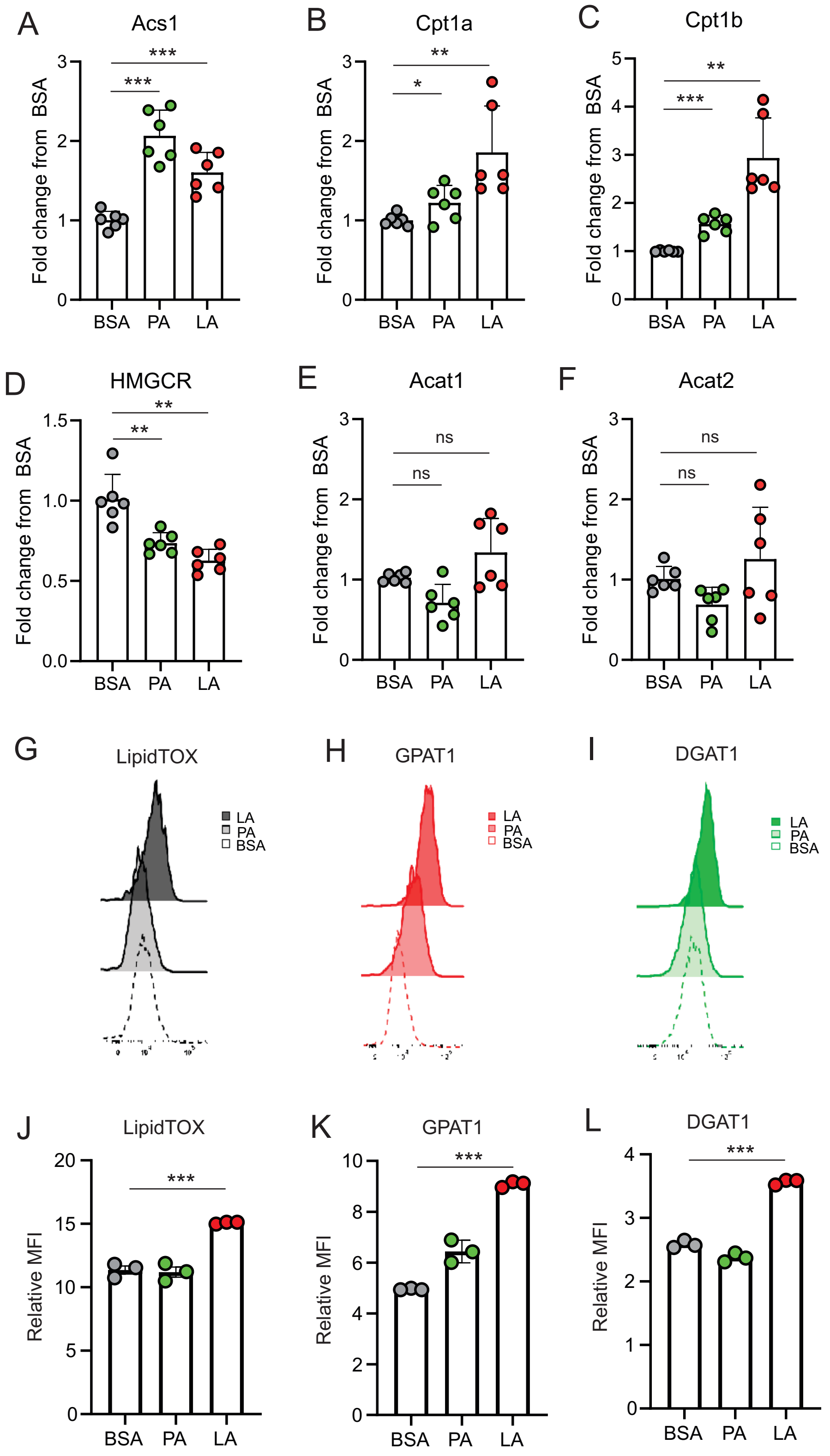
A
Acs1
Fold change from BSA
BSA PA LA
B
Cpt1a
C
Cpt1b
D
HMGCR
E
Acat1
F
Acat2
G
LipidTOX
LA
PA
BSA
H
GPAT1
I
DGAT1
J
LipidTOX
Relative MFI
K
GPAT1
L
DGAT1

**Supplementary Figure 2 LA induces key enzyme expression in triglyceride biosynthesis.**

A-F, Analysis of FA metabolism-related genes, including Acs1 (A), Cpt1a (B), Cpt1b (C), HMGCR (D), Acat1 (E), Acat2 (F) in macrophages treated with BSA, PA or LA (400μM) for 4h.

G-I, Flow cytometric staining for LipidTOX (G), GPAT1 (H), and DGAT1 (I) expression in macrophages treated with BSA, PA or LA (400 μM).

J-L, Quantification of LipiTOX (J), GPAT1 (K), and DGAT1 (L) expression by mean fluorescent intensity (MFI) in macrophages treated with BSA, PA or LA (400 μM).

Data are shown as mean ± SD in panel A-F and J-L (* $p \le 0.05$, ** $p \le 0.01$, **** $p \le 0.0001$, ns, non-significant, unpaired Student t test).

### S-Figure 3

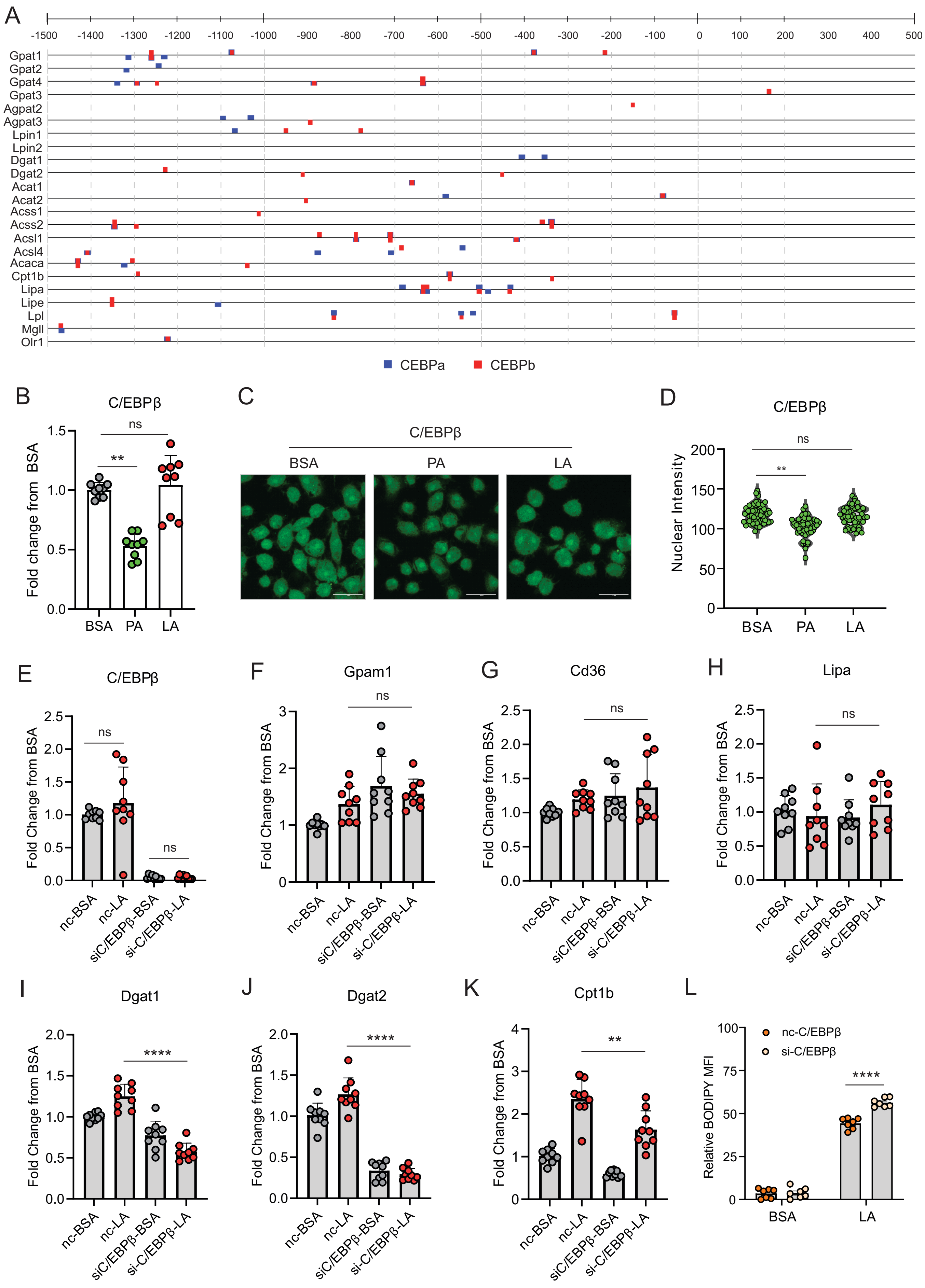
A
-1500 -1400 -1300 -1200 -1100 -1000 -900 -800 -700 -600 -500 -400 -300 -200 -100 0 100 200 300 400 500
Gpat1
Gpat2
Gpat4
Gpat3
Agpat2
Agpat3
Lpin1
Lpin2
Dgat1
Dgat2
Acat1
Acat2
Acss1
Acss2
Acsl1
Acsl4
Acaca
Cpt1b
Lipa
Lipe
Lpl
Mgll
Olr1
CEBPa
CEBPb
B
C/EBPβ
Fold change from BSA
BSA PA LA
C
C/EBPβ
BSA PA LA
D
C/EBPβ
Nuclear Intensity
BSA PA LA
E
C/EBPβ
F
Gpam1
G
Cd36
H
Lipa
Fold Change from BSA
nc-BSA
nc-LA
siC/EBPβ-BSA
si-C/EBPβ-LA
I
Dgat1
J
Dgat2
K
Cpt1b
L
nc-C/EBPβ
si-C/EBPβ
Relative BODIPY MFI
BSA LA
ns
**
****

**Supplementary Figure 3 LA-induced lipid accumulation in macrophages was undependable on CEBPβ**

A, CiiiDER software package was used to identify potential C/EBPα and C/EBPβ transcription factor binding sites in regulatory regions of relevant lipid droplet and lipolysis genes.

B, Real-time PCR analysis of CEBPβ expression macrophages treated with BSA, PA and LA for 4 hours.

C,D, Confocal analysis for CEBPβ staining (green) in macrophages treated with BSA, PA and LA. Quantification of CEBPβ staining is shown in panel D.

E-K, Macrophages was transfected with CEBPβ or control siRNA, and then treated with BSA or LA for 4 hours. Expression of CEBPβ (E), lipid droplet formation and other FA-metabolism related genes, including Gpam1 (F), Cd36 (G), Lipa (H), Dgat1 (I), Dgat2 (J), Cpt1b (K) was determined by real-time PCR analysis.

L, Measurement of lipid droplet formation by BODIPY staining in CEBPβ-silencing macrophages treated with BSA or LA.

Data are shown as mean ± SD in panel L-O (** $p \leq 0.01$, **** $p \leq 0.0001$, ns, non-significant, unpaired Student t test).

### S-Figure 4

A

| CEBPA Correlated Gene | Spearman's Correlation | p-Value | q-Value |
|---|---|---|---|
| FABP4 | 0.35 | 1.26E-32 | 7.68E-31 |
| FABP3 | 0.226 | 5.55E-14 | 3.84E-13 |
| FABP5 | 0.221 | 1.78E-13 | 1.16E-12 |
| FABP9 | 0.129 | 0.00002077 | 0.00005225 |
| FABP5P3 | 0.08 | 0.008505 | 0.0148 |
| FABP6 | 0.0768 | 0.0115 | 0.0196 |
| FABP12 | -0.0628 | 0.0389 | 0.0599 |
| FABP2 | 0.034 | 0.264 | 0.332 |
| FABP7 | 0.0249 | 0.414 | 0.488 |
| FABP1 | 0.006119 | 0.841 | 0.874 |

B

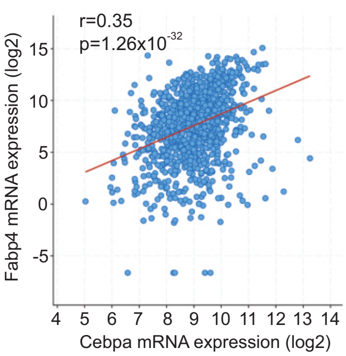


C

| FABP4 Correlated Gene | Spearman's Correlation | p-Value | q-Value |
|---|---|---|---|
| CEBPA | 0.35 | 1.26E-32 | 3.09E-31 |
| CEBPG | -0.238 | 1.89E-15 | 1.69E-14 |
| CEBPZ | -0.22 | 2.29E-13 | 1.71E-12 |
| CEBPD | 0.219 | 3.01E-13 | 2.22E-12 |
| CEBPE | 0.161 | 1.11E-07 | 4.58E-07 |
| CEBPA-DT | 0.151 | 5.76E-07 | 2.17E-06 |
| CEBPB | 2.74E-03 | 0.928 | 0.946 |

D

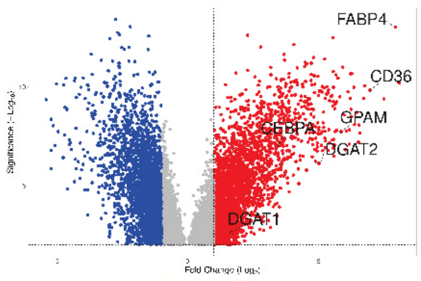


VolcaNoseR - GSE42568

| Name | Change | Fold change (log2) | Significance |
|---|---|---|---|
| FABP4 | Increased | 7.918 | 13.027 |
| CD36 | Increased | 6.941 | 9.828 |
| GPAM | Increased | 5.854 | 7.739 |
| DGAT2 | Increased | 5.013 | 6.119 |
| CEBPA | Increased | 3.43 | 6.996 |
| DGAT1 | Increased | 1.684 | 2.6 |

E

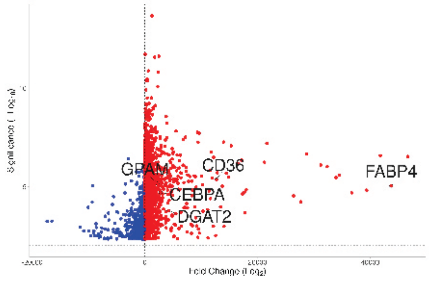


VolcaNoseR - GSE21422

| Name | Change | Fold change (log2) | Significance |
|---|---|---|---|
| FABP4 | Increased | 43690.227 | 5.038 |
| CD36 | Increased | 12712.72 | 5.353 |
| DGAT2 | Increased | 4197.407 | 3.772 |
| CEBPA | Increased | 2796.453 | 4.699 |
| GPAM | Increased | 2208.953 | 5.153 |

F

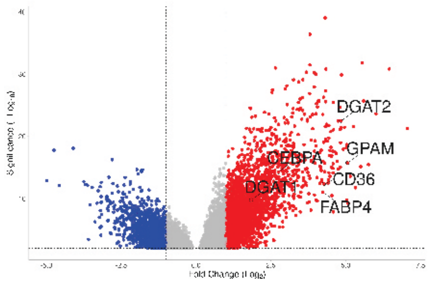


VolcaNoseR - GSE29431

| Name | Change | Fold change (log2) | Significance |
|---|---|---|---|
| GPAM | Increased | 5.061 | 15.741 |
| DGAT2 | Increased | 4.84 | 22.475 |
| CD36 | Increased | 4.274 | 12.363 |
| FABP4 | Increased | 4.234 | 11.024 |
| CEBPA | Increased | 2.754 | 14.231 |
| DGAT1 | Increased | 1.842 | 9.784 |

G

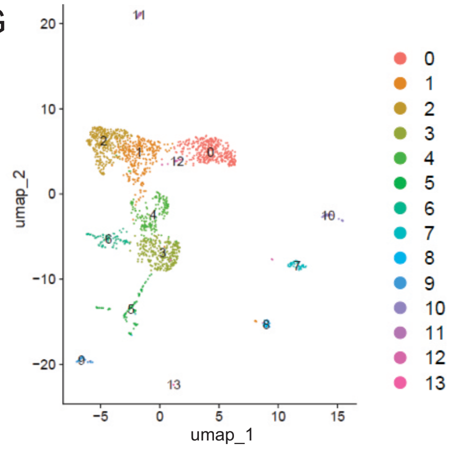


H

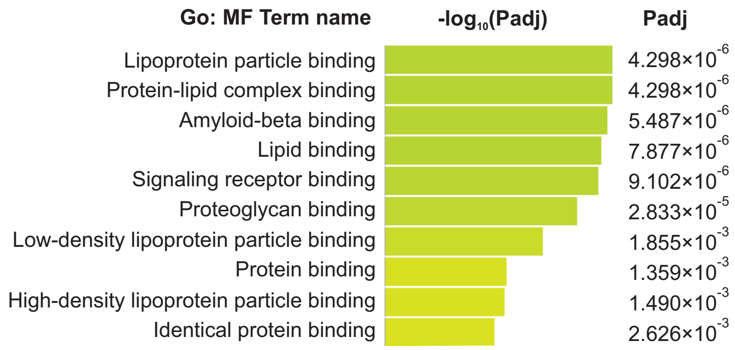


I J K L

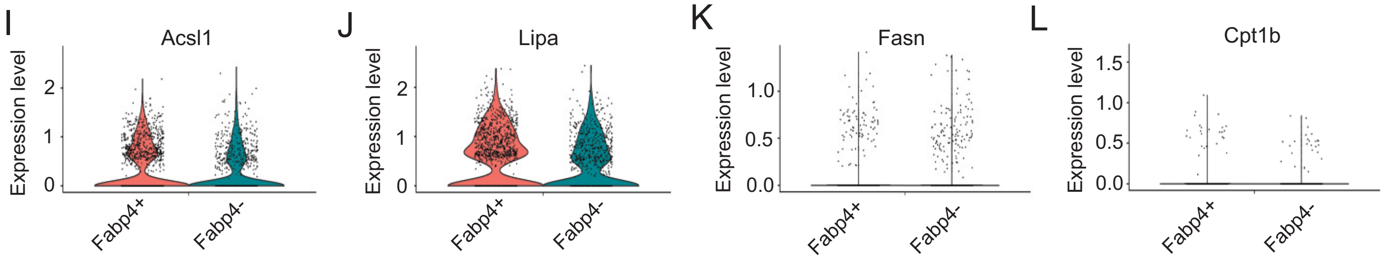


M N O P

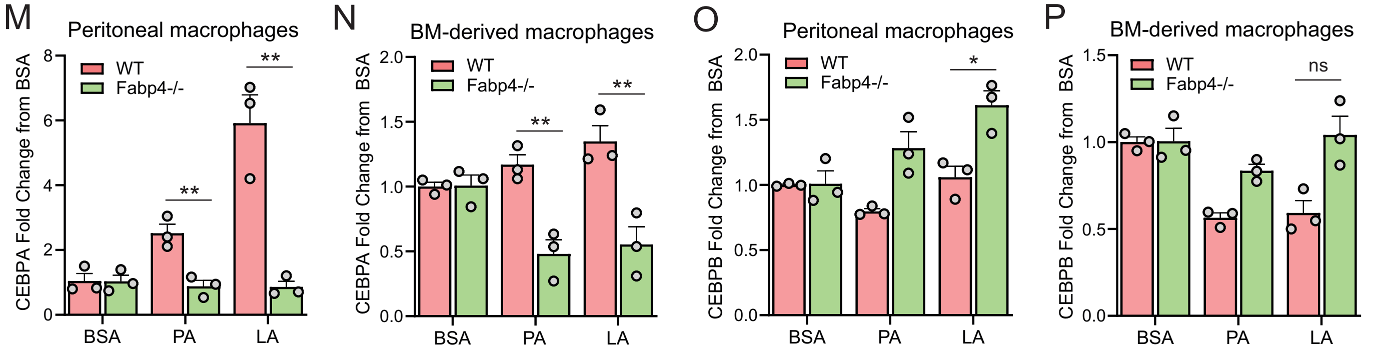

**Supplementary Figure 4 FABP4 mediates LA-induced CEBPα, but not CEBPβ, activation**

A,B, Spearman's correlation analysis of CEBPα with FABP family members, especially FABP4 with CEBPα (B), using cBioportal breast cancer TCGA database.

C, Spearman's correlation analysis of FABP4 with CEBP family members using cBioportal breast cancer TCGA database.

D-F, Differential gene expression analysis of FABP4, GPAM, DGAT1, DGAT2, CEBPA, and CD36 in published breast cancer microarray databases, including GSE42568 (D), GSE21422 (E) and GSE29431 (F).

G, UMAP of different subsets of splenic macrophages using single-cell RNA sequence analysis

H, Molecular function pathway analysis of the integrated conserved genes in the FABP4-positive clusters by g:Profiler.

I-L, Violin plots showing relative expression levels of genes, including Acs1(I), Lipa (J) and Fasn (K) and Cpt1b (L) between FABP4+ vs FABP4- macrophages in the splenic macrophage single cell RNA sequencing analysis.

M, Measurement of CEBPA gene levels in WT and FABP4$^{-/-}$ peritoneal macrophages treated with BSA, PA, or LA, respectively, for 4 hours.

N, Measurement of CEBPA gene levels in WT and FABP4$^{-/-}$ bone marrow derived macrophages treated with BSA, PA, or LA, respectively, for 4 hours.

O, Measurement of CEBPB gene levels in WT and FABP4$^{-/-}$ peritoneal macrophages treated with BSA, PA, or LA, respectively, for 4 hours.

P, Measurement of CEBPB gene levels in WT and FABP4$^{-/-}$ bone marrow derived macrophages treated with BSA, PA, or LA, respectively, for 4 hours.

Data are shown as mean ± SD in panel I-P (p* $p \leq 0.05$, ** $p \leq 0.01$, ns, non-significant, unpaired Student t test).

### S-Figure 5

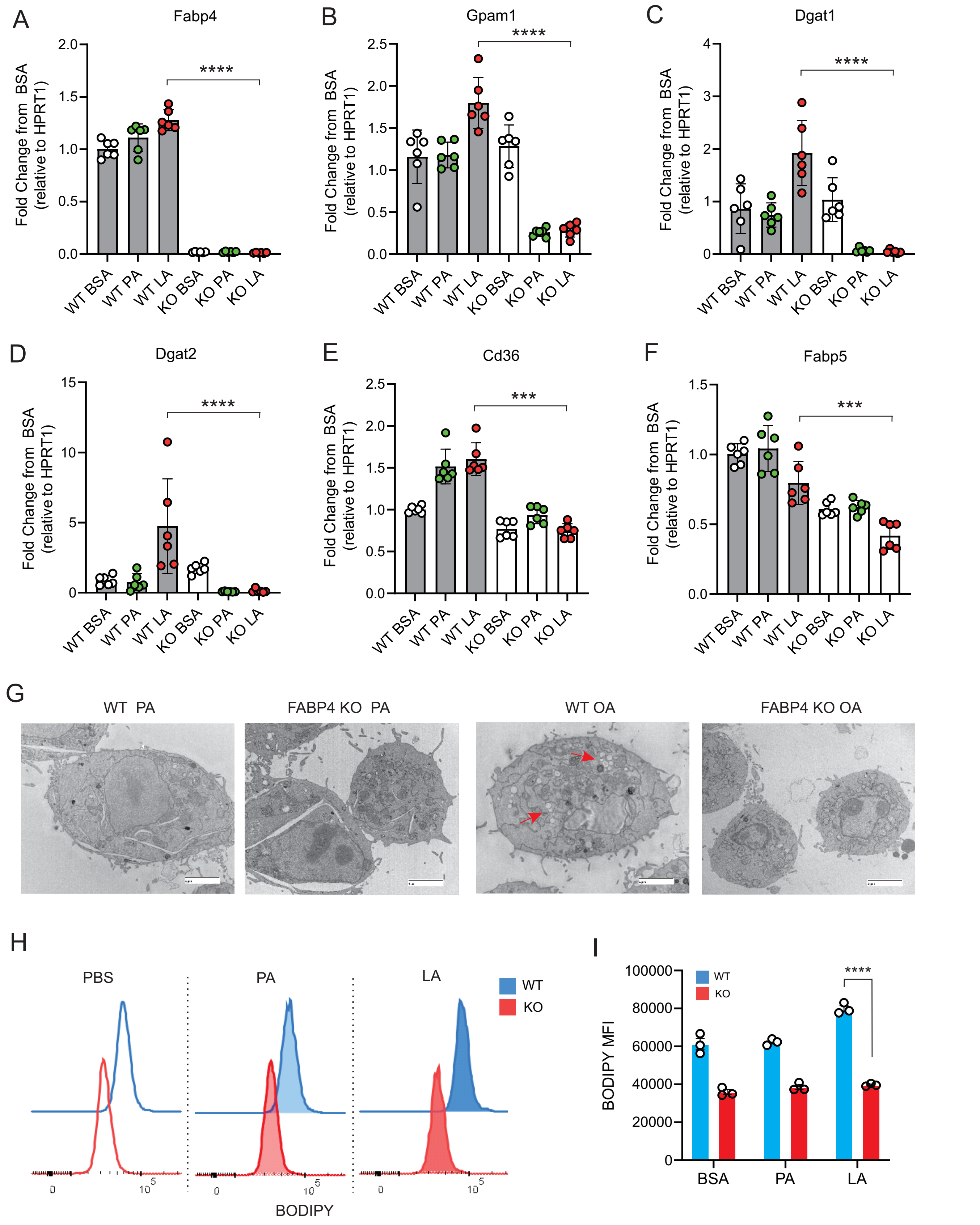
A
Fabp4
B
Gpam1
C
Dgat1
D
Dgat2
E
Cd36
F
Fabp5
Fold Change from BSA
(relative to HPRT1)
****
***
WT BSA
WT PA
WT LA
KO BSA
KO PA
KO LA
G
WT PA
FABP4 KO PA
WT OA
FABP4 KO OA
H
PBS
PA
LA
WT
KO
BODIPY
I
BODIPY MFI
BSA
PA
LA

**Supplementary Figure 5 Deficiency of FABP4 reduces lipid droplet formation in macrophages**

A-F, Real-time PCR analysis of FABP4 (A) and genes encoding key enzymes for triglycerol biosynthesis, including Gpam1 (B), Dgat1 (C), Dgat2 (D), Cd36 (E), Fabp5 (F) in WT and FABP4 KO peritoneal macrophages treated with BSA, PA and LA (400μM).

G, Transmission electron microscope showing lipid droplet formation (red arrows) in WT and KO macrophages treated with either PA or OA for 4 hours.

H, I, Flow cytometric analysis of BODIPY staining (G) and mean fluorescent intensity (MFI) in peritoneal macrophages treated with BSA, PA, or LA (400μM).

Data are shown as mean ± SD (*** $p \le 0.001$, *** $p \le 0.001$, unpaired Student t test).

### S-Figure 6

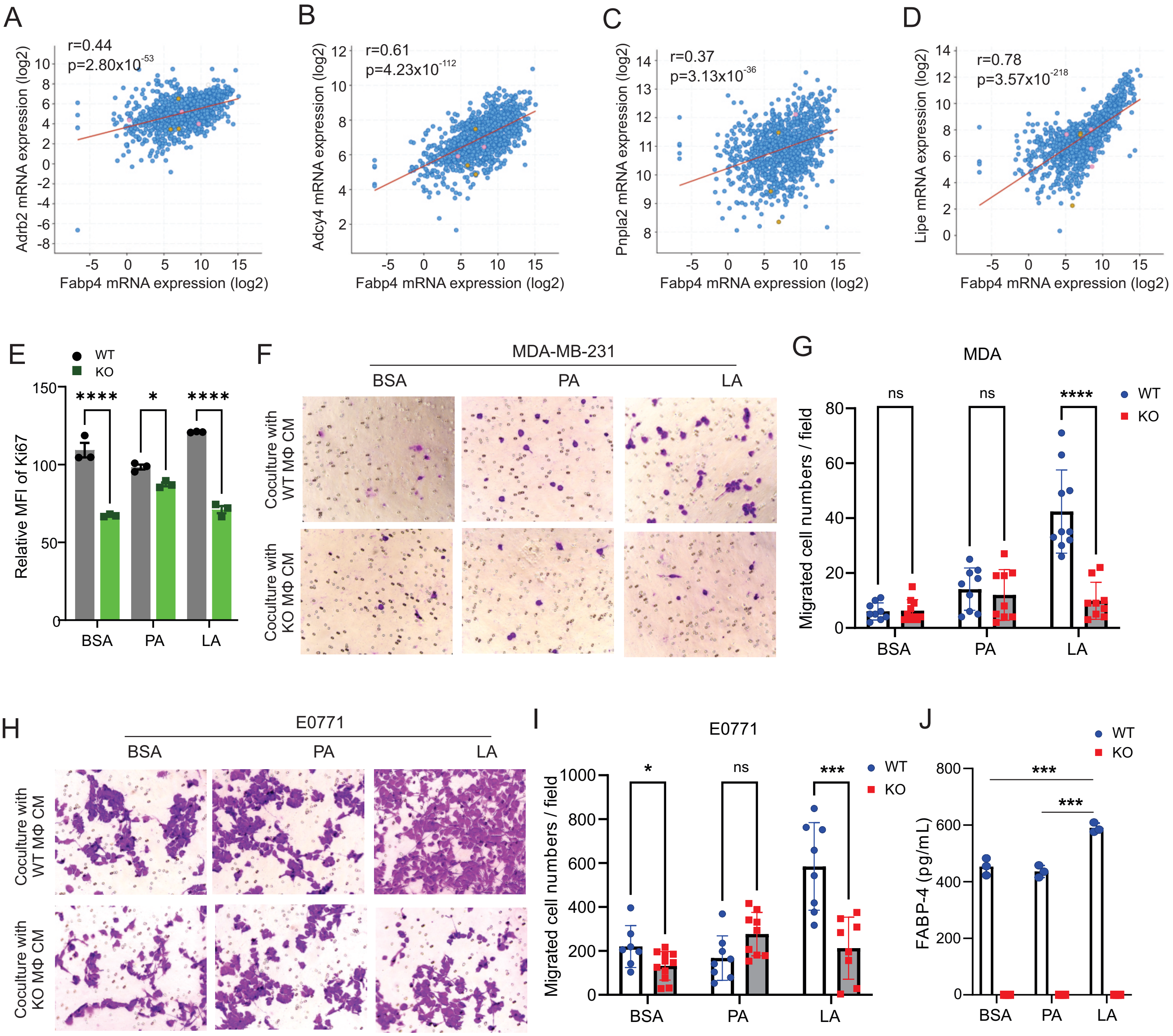

A
r=0.44
p=2.80x10-53
Adrb2 mRNA expression (log2)
Fabp4 mRNA expression (log2)
B
r=0.61
p=4.23x10-112
Adcy4 mRNA expression (log2)
Fabp4 mRNA expression (log2)
C
r=0.37
p=3.13x10-36
Pnpla2 mRNA expression (log2)
Fabp4 mRNA expression (log2)
D
r=0.78
p=3.57x10-218
Lipe mRNA expression (log2)
Fabp4 mRNA expression (log2)
E
WT
KO
Relative MFI of Ki67
BSA
PA
LA
F
MDA-MB-231
BSA
PA
LA
Coculture with WT MΦ CM
Coculture with KO MΦ CM
G
MDA
Migrated cell numbers / field
ns
ns
****
WT
KO
BSA
PA
LA
H
E0771
BSA
PA
LA
Coculture with WT MΦ CM
Coculture with KO MΦ CM
I
E0771
Migrated cell numbers / field
*
ns
***
WT
KO
BSA
PA
LA
J
WT
KO
FABP-4 (pg/mL)
***
***
BSA
PA
LA

**Supplementary Figure 6 FABP4 promotes lipolysis and tumor migration**

A-D, Spearman's correlation analysis between FABP4 and key genes mediating lipolysis, including Adrb2 (A), Adcy4 (B), Pnpla2 (C), and Lipe (D) using the cBioportal breast cancer TCGA database.

E, Measurement of cell proliferation marker Ki67 in E0771 tumor cells after coculture with FA-treated FABP4 WT or KO macrophages for 24 hours.

F,G, After treatment with BSA, PA, or LA (400uM) for 4 hours, FABP4 WT and KO macrophages were cultured in FBS-free RPMI-1640 for 24h. The cultural conditional medium was collected for tumor migration assays. MDA-MB-231 migration and quantification are shown in panel F and G, respectively.

H, I, E0771 tumor migration assays were performed using the conditional medium collected as above plus 1% FBS. Tumor migration and quantification are shown in panel H and I, respectively.

J, ELSIA measurement of FABP4 levels in macrophage/tumor coculture medium.

Data are shown as mean ± SD in panel G, I and J (p* $p \leq 0.05$, *** $p \leq 0.001$, ns, non-significant, unpaired Student t test).

Cd163 gene expression
cohort divided at median of gene expression

### S-Figure 7

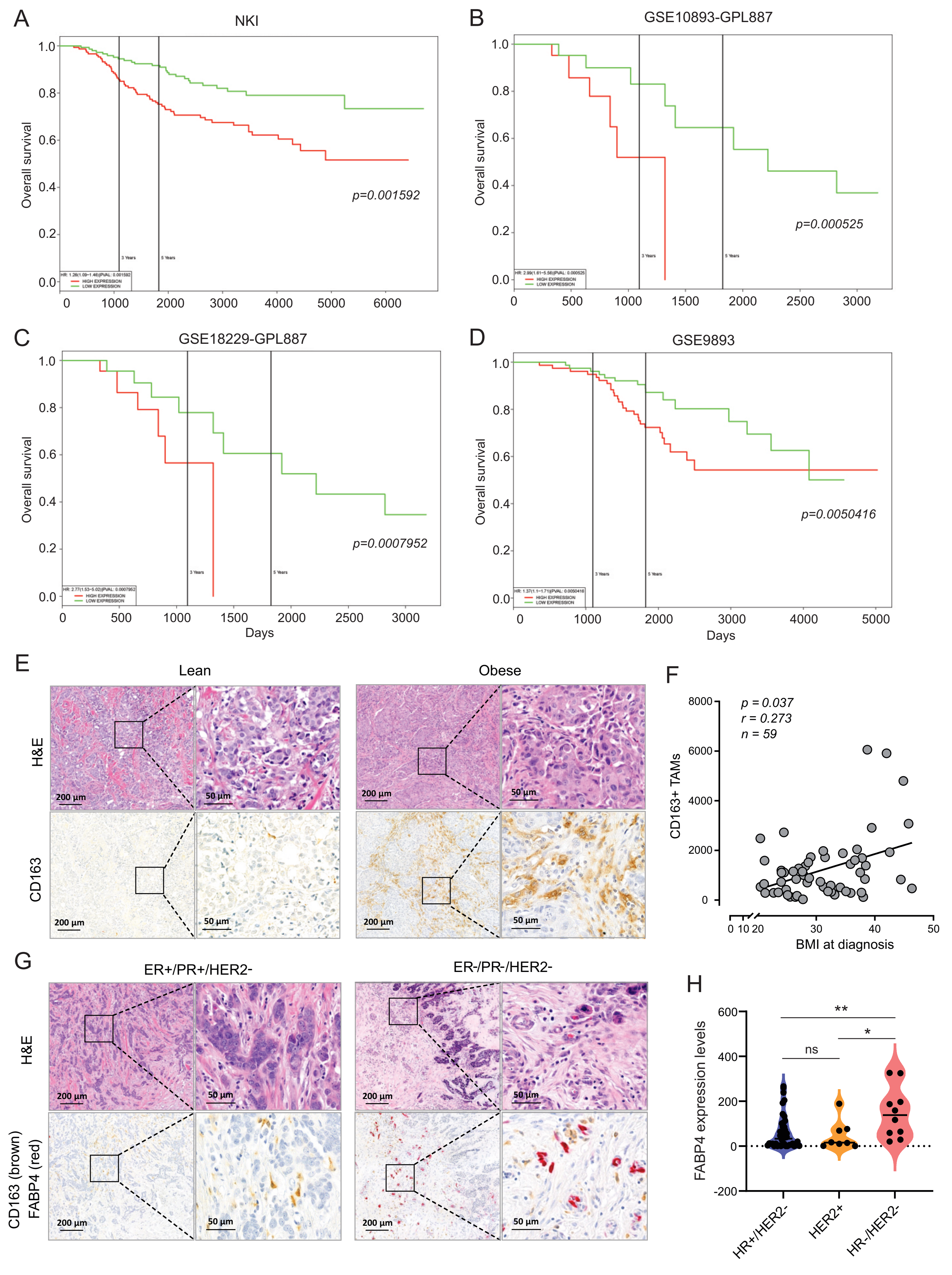

**Supplementary Figure 7 Association of TAMs with obesity and survival of breast cancer patients**

A-D, PROGgene analysis of the association between CD163 expression levels and breast cancer survival using various publicly accessible breast cancer databases, including NKI (A), GSE10893 (B), GSE18229 (C) and GSE9893 (D).

E, Comparison of H&E and CD163 staining (brown) between lean and obese breast cancer tumors in breast cancer patients.

F, Spearman’s correlation analysis between CD163 expression in breast cancer tissues and BMI at diagnosis of breast cancer patients.

G, H&E and immunohistochemistry staining of FABP4 (red) in ER+ and triple negative breast cancer tissues.

H, Analysis of FABP4 expression in hormone receptor+/HER2-, HER2+ and triple negative breast cancers.

Data are shown as mean ± SD in panel H (p* $p \leq 0.05$, ** $p \leq 0.01$, ns, non-significant, unpaired Student t test).
